# Optimizing root measurements in rhizotrons

**DOI:** 10.1101/587329

**Authors:** NK Ytting, JA Kirkegaard, K Thorup-Kristensen

## Abstract

**Background and Aims:** The line intersect method is widely used in rhizotron and minirhizotron studies to quantify roots and study cultivar and treatment differences in root growth. We investigated ways to optimize the line intersect method and root depth measurements with respect to data variability and the time spent on counting roots.

**Methods:** Root intensity was measured with three different grid patterns and different lengths of counting line on 2 m long transparent tube rhizotrons. Rooting depth was recorded by measuring the depth of the deepest root and by measuring the depth below which 5, 10 and 25 roots were observed.

**Results:** For root intensity the coefficient of variation (CV) was reduced 10-50 percentage points for grids that distributed counting lines equally across the measured area compared to using a restricted centralized area. In addition, the CV approached an asymptote of around 40 % when more than 50 root intersections per grid were observed. Further we show that recordings of the deepest root gave the most variance and least difference between means with a p-value of 0.65 for difference between cultivars. In contrast, a significant difference between cultivar rooting depths (p = 0.01) was found when using the depth below which 25 roots were observed.

**Conclusion:** We propose the use of grid designs adapted to different root densities to decrease time spent on counting roots at high root intensities, and minimize data variability at low root intensities. Further on rooting depth measurements including more roots may be a more useful parameter statistically to reveal variety or treatment differences in rooting depth.

## Introduction

Humanity faces a major challenge in coming decades to provide both food and biofuels for the rapidly growing population with diminishing water and nutrient resources (Foley et al., 2011). To meet the challenge plant scientists have focused on ways to increase crop productivity and efficiency, and manipulating plant root systems is considered by some to be the key (Lynch, 2007; Lynch and Brown, 2012). As a result there is increasing interest in the study of root growth and function in soil and an increasing inclusion of root phenes as selection parameters in the development of new crop genotypes (Lynch, 2007; Wasson et al., 2014). However, progress remains limited as suitable methodologies for root studies remain labor intensive and indirect, as roots are hidden in the soil. Several recent studies have focused on developing faster and more automated methods for root studies. These methods have mainly focused on techniques where roots of very young plants are studied under artificial conditions in the lab (Yazdanbakhsh and Fisahn, 2009; Iyer-Pascuzzi et al., 2010; Topp et al., 2013), and are far removed from the field conditions where roots grow and function. The ultimate goal is to study and understand root growth and function in the field (Gregory et al., 2009), so more efficient methods for root studies of older and larger plants growing in soil media is required.

For root studies of soil-grown plants, non-destructive methods are desirable because root excavation methods are both time and labour intensive. Methods such as rhizotrons and minirhizotrons offer possibilities for non-destructive measurements of the dynamic processes of root growth and traits such as rooting depths and intensities. Rhizotrons were originally covered underground walkways with glass walls that allowed for direct observations of root growth (Taylor et al., 1990). Later, growth containers of any kind with transparent sides, including transparent tubes, have been referred to as rhizotrons (Mattina et al., 2004; Nagel et al., 2012), but the terms tube rhizotrons (Ytting et al., 2014), rhizobox (Watt et al., 2006) and root observation boxes (Thaler and Pagés, 1996) have all been used to describe these containers. Minirhizotrons are long, often > 1 m in length, glass tubes inserted in the soil, that allow for direct imaging of root systems (Johnson et al., 2001). In both types of installations, images of the soil-rhizotron interface can be taken at regular spatial and time point intervals.

Many software solutions have been developed to provide information on root architectural traits from root images. Only a few of these solutions are applicable for images of roots with soil as a background and none of them are fully automatized for discrimination between roots and soil (Lobet et al., 2013). The available software requires manual tracking of roots by clicking along the individual root with the curser or a finger. This work step can take thousands of hours per experiment (Zeng et al., 2010). Therefore, the line intersect method first described by Newman (1966) is still a widely used method for measuring root intensities in different soil layers of rhizotrons and minirhizotrons (Thorup-Kristensen, 2001; Kristensen and Thorup-Kristensen, 2004; Christiansen et al., 2006; Kristensen and Thorup-Kristensen, 2007; Ulas et al., 2012; Wang et al., 2014; Andersen et al., 2014; Ytting et al., 2014). The method was originally designed to estimate total length of roots from a sample of roots washed free from soil. Later, the method was modified by March (1971) and validated by Tennant (1975). When using the line intersect method on rhizotrons or minirhizotrons, a grid of counting lines is superimposed on the soil-rhizotron interface and root-line crossings are counted either directly, or in photographs which are compiled and counted later. Often the method is used to give information on root intensity in the measured area with the unit of root intersections per m counting line. The unit is hereby independent of the area of measurement and length of counting line.

There are two challenges when using the line intersect method. Firstly, the time required for root counting is often the major bottleneck for data acquisition, and secondly there is often high data variability. The focus of this study was to investigate strategies to decrease the time spent on counting roots without compromising accuracy.

One approach to decrease the time spent on counting roots when using the line intersect method is to decrease the length of counting line, but this comes with the potential cost of increased data variance. Root growth happens in predetermined patterns, as the growth direction of main roots is gravity controlled with lateral roots appearing along the axis of the growing main roots. Pores and cracks offer channels of reduced physical resistance leading to increased root growth compared to the surrounding soil and often several roots can be observed in a single soil pore (Stirzaker et al., 1996; White and Kirkegaard, 2010). As a consequence, roots are not always evenly distributed in the soil and spatial autocorrelation can be an issue of concern. We hypothesized that the grid designs are of major importance for obtaining observations representative for the total observation area. The length of analyzed counting line can be reduced in different ways. One way is to reduce the observation area to a sub-area while maintaining the size of grid elements (design A). By doing so, a smaller area of observation is needed which reduces the number of images needed. An alternative method is to keep the total observation area but reduce the length of counting line by increasing the size of grid elements (design B) or by distributing smaller pieces of discontinuous grid lines across the studied area (design C) (Fig. 1).

**Figure 1:**
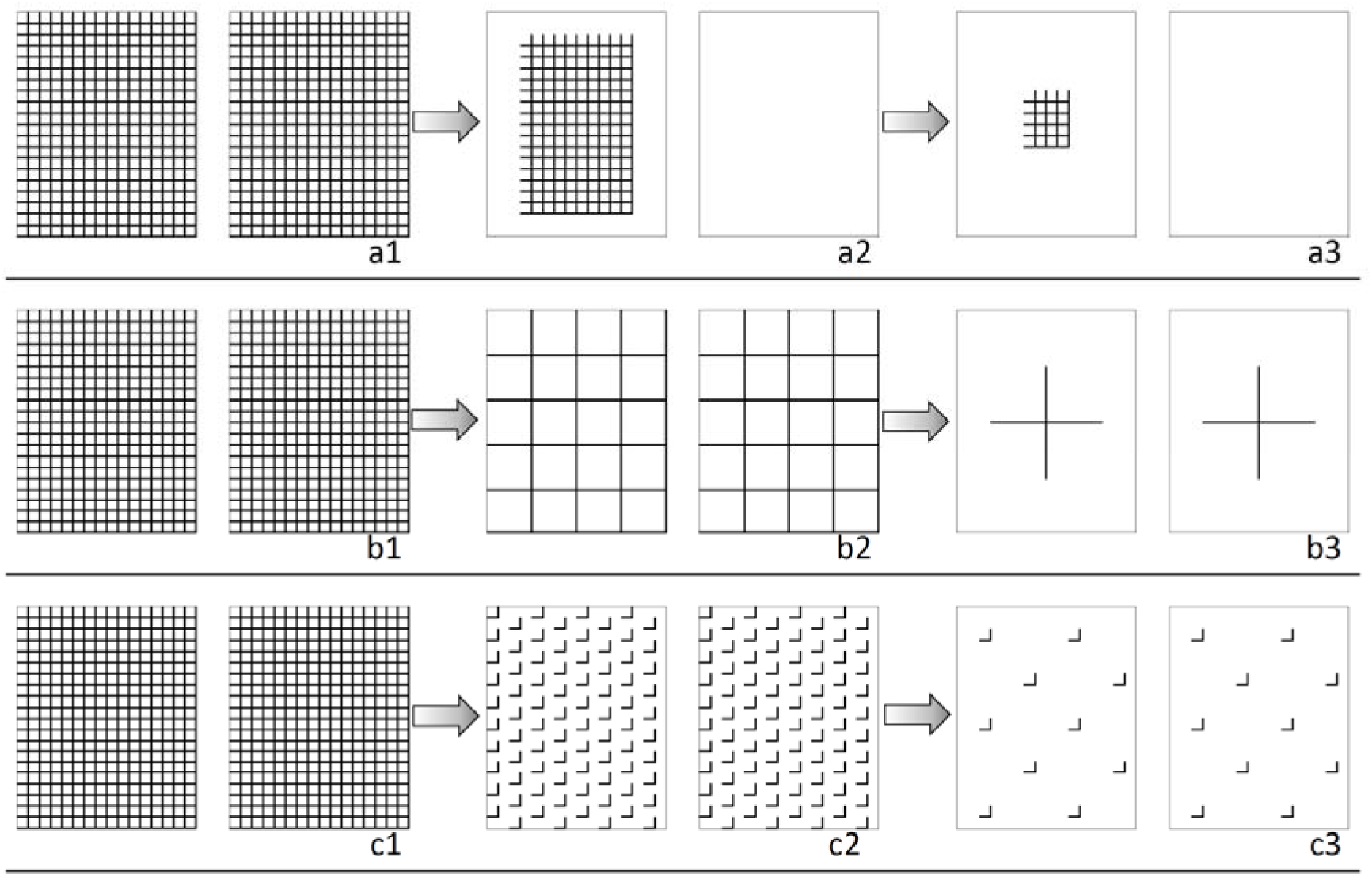
Grids superimposed on pictures of transparent tube rhizotrons. On each tube rhizotrons one grid was placed on the front side and one grid on the back side. Dimensions of each grid were 80 mm × 100 mm. Total lengths of counting lines are 6.4 m (a1,b1 and c1), 1.6 m (a2,b2 and c2) and 0.2 m (a3,b3 and c3). Grid element size of 5 × 5 mm on entire measuring area (a1,b1 and c1). Examples of different approaches (design A, B and C) to decrease the length of counting line in a defined measuring area on the tube rhizotrons. In design A grid element size is maintained and measuring area decreased (a2 and a3). In design B grid element size is increased and measuring area maintained (b2 and b3). In design C the number of small line pieces is decreased and measuring area maintained. Small lines pieces equally scattered on measuring area (c2 and c3).

In addition we hypothesize that data variance is negatively correlated to total root counts per grid – that is when root observations per measured area decreases, the variance will increase. This is because the outliers are given more weight when few observations contribute to the mean and variance (Dean and Dixon, 1951). With few visible roots in the observation area a higher density of grid lines is needed for maintaining low variance in data. Rather than defining a fixed pattern suitable for all root intensities, we propose to define the minimum number of root intersections to count per measuring area based on root intensity. Thus, by modifying the grid to match the root intensity, both data variance and counting time can be reduced. We imagine that this can be done by using a grid with grid lines of different colors; grid lines of each color are equally distributed across the measuring area and each color has the same total length. If the minimum number of root intersections is not met with the first color of lines, the next color of lines is included in the counting and calculation of root intensity, and so forth for the rest of the colors.

Root depth measurements of soil-grown plants have been carried out in a range of ways in the literature depending on the scope of the study. The term “rooting depth” has no clear definition, and often it is not the maximum root depth, but the effective rooting depth of the root system that is of interest. Individual roots can be much deeper than average maximum root depth (Kirkegaard and Lilley, 2007), but without much function for water and nutrient uptake. The term “rooting depth” has been variously used to describe 1) average maximum root depth, measured as the average depth of the deepest roots observed (Thorup-Kristensen, 2001; Svoboda and Haberle, 2006; Kirkegaard and Lilley, 2007; Acuña and Wade, 2012; Wasson et al., 2014; Ytting et al., 2014), 2) the average depth, below which 3 roots were observed (Thorup-Kristensen, 1998), 3) the average depth, below which 25 roots were observed (Thorup-Kristensen, 1998). The deepest observed root will often not be the absolute deepest root of the plants or crop stands. In studies using core methods, minirhizotrons and rhizotrons, the rooting depth is both a measure of root length density (RLD) in depth and rooting depth. Because of the low RLD in the deepest part of the root system, the likelihood of observing or finding the depth of the deepest roots of individual plants or crop stands is correlated to the volume of soil that is examined. An increase in the observation area would result in a decrease of the coefficient of variation (Ping et al., 2010) but at the same time, significantly increase resource demand.

For rooting depth measurements we hypothesize that measurements of the deepest roots have higher variability than measurements of the depth below which a pre-determined number of roots are observed. This is due to the often increased RLD at decreasing root system depth. Smaller differences in rooting depths among genotypes or treatments can be measured if recordings of the depth below which a larger number of roots (e.g. 25 roots) are observed and used instead of recordings of the deepest root. Given a certain minimum root density is generally required for functionality in terms of water or nutrient uptake (Noordwijk, 1983) and others), estimations based on a larger number of roots may also have more functional meaning.

The focus of this study was to optimize the line intersect method and root depth measurements in rhizotron and minirhizotron studies. We used studies of wheat plants growing in 2 m deep tube-rhizotrons to investigate improvements in the time required and efficiency of root measurements without compromising accuracy. This was achieved by comparing CVs for different length of counting lines and grid patterns and by adjusting root depth measurements according to the average depth below which a predefined number of roots were observed.

## Materials and Methods

The experiment was conducted from May to July 2012 under a glass-roof with open sides in a field in Taastrup, Zealand, Denmark 55°40’90.35”N and 12°18’24.84E. The glass roof transmitted a minimum of 50 % of the photosynthetic active radiation depending on the height of the sun and amount of diffuse light. The experiment in which the methodologies for root counting were compared was a complete randomized block design with four N treatments, six wheat cultivars and four replicates.

### Tube rhizotrons

The experimental tube rhizotrons were made of transparent PVC tubes (OTV Plast A/S) filled with soil. The tubes were 74 mm in inner diameter and 80 mm in outer diameter and consisted of a lower (0.5 m) and an upper (1.5 m) section. The lower section was halved lengthwise and taped together for ease of access to the soil column for later root washing. The combined height of the tube sections was 2.0 m. The bottom of the tubes were covered with plastic net (1 × 1 mm) fixed with tape to retain the soil within the column while allowing drainage. To mimic the field soil profile, the soil used to fill the tubes was excavated subsoil (0.3 - 0.6 m) and topsoil (surface 0 - 0.3 m) from a field at the experimental farm. The soils were separately excavated, air dried, sieved on a 10 mm sieve and mixed thoroughly in a concrete mixer prior to filling of the tubes.

The subsoil was filled into the bottom of the tube rhizotrons and topsoil in the top 0.25 m. The subsoil was a sandy soil, and the topsoil a loamy sand; soil characteristics were described in Ytting et al., (2014). For the lower tube sections only, the subsoil was prepared with four levels of nitrogen (ammonium nitrate) (0, 2.5, 5 and 10 mg N kg-1 soil) and supplemented with other macro and micro nutrients at a standard rate before being packed into the tubes.

The subsoil was compacted by adding it loosely into the upright tubes while they were held on a vibrating plate powered by a NTK 25 AL piston oscillator (NTK® Oscillators) set at 6 bar and an amplitude of 5.8 mm. Topsoil (1.4 kg tube-1) was subsequently added loosely in the top 0.25 m of the tube and compacted by hand to a set height of 15 mm below the top edge of the tube.

The differences in mean bulk density among tube rhizotrons did not exceed ± 0.6 %. The subsoil in the upper tube sections was packed at a gravimetric water content of 4.6 % and the final mean dry bulk density was 1.52 g cm-^3^. The topsoil was packed at 3 % gravimetric moisture and the mean dry bulk density was 1.44 g cm-^3^. The subsoil in lower tube sections was packed with a gravimetric water content of 15 % and mean bulk density of 1.67 g cm-^3^.

### Frames

The tube rhizotrons were fixed on wooden frames, each holding 20 tube rhizotrons. The frames were insulated on all sides and on the top by a box (2.0 m high × 1.2 m × 0.8 m) made of RIALET® Foamalux 10 mm plates (Brett Martin Ltd, UK), a white foamed PVC product. Holes were made in the top for the tube rhizotron to go through the insolating layer. The soil surface and above ground parts of the plants were exposed to the environment while the rest of the tube rhizotrons were enclosed within the box. Each tube rhizotron was further wrapped individually in corrugated cardboard to exclude all light.

### Seeding, management and harvest

Each tube rhizotron was seeded with four wheat seeds (*Triticum aestivum* L.) on 19 May. The six different wheat cultivars (three Australian spring types and three North European winter types) were chosen based on earlier reported differences in root traits. Following emergence, the plants were thinned to one plant in each tube rhizotron, corresponding to a plant density of 232 plants m-2 soil surface area.

The tube rhizotrons were irrigated individually from the top using a drip water dispenser of the type Iriso enkeltdryppere (Modious ApS, Denmark) in order to maintain moisture content within 50-80 % of field capacity. Fungal diseases was observed once and subsequently treated with the fungicide.

The shoots were harvested on 26 July at developmental stage 53 to 69 for spring types and 29 for winter types (BBCH scale) by cutting below the stem base at the soil surface. After harvest, the upper 1.5 m tube rhizotrons sections were separated from the lower 0.5 m tube sections. The lower 0.5 m sections were stored at 4°C prior to further root measurements.

### Root measurements

For some of the root measurements, the number of cultivars and nitrogen (N) treatments was reduced (Table 1). One cultivar, with unusually poor germination and growth, presumably due to poor seed quality, was excluded from analysis. For the analysis of grid patterns and length of observation line only three cultivars and two N treatments were included. This was done to minimize the substantial processing time of the pictures, and fewer observations were sufficient to answer this research question. For the same reason, a reduced number of tubes were included in the analysis of cultivar effects on root intensity.

**Table 1:** For each type of root measurements this table shows the number of cultivars, nitrogen treatments and total number of tubes used.

| Type of measurement | Number of cultivars | Number of N treatments | n | Figure |
| --- | --- | --- | --- | --- |
| Correlation between root density of washed roots and root intensity measured with the line intersect method | 6 | 4 | 96 | 2 |
| Root intensity for analysis of grid patterns and reduction of counting line | 3 | 2 | 24 | 3 |
| Rooting depth | 5 | 4 | 80 | 4 |
| Cultivar and nitrogen effects on root intensity | 3 | 2 | 24 |  |

### Root intensity measurements on tube rhizotrons

On 25 July - the final day of the experiment - the root intensity in the 1.71 – 1.79 m layer was measured by superimposing grids on the transparent tube rhizotrons and counting root-line intersections. The grid size for this measurement was 60 × 80 mm with a grid element size of 20 × 20 mm and a total length of counting line of 0.48 m. Root intensity measurements with the line intersect method have been used before, however descriptions of root recording details are scarce (e.g. Thorup-Kristensen 2001, Christiansen et al. 2006 and Ulas et al. 2012). A clearer approach is needed to establish when a root is considered to be intersecting a line. Newman (, 1966) originally defined this by only counting an intersection if the counting line crossed the center line of the root. Tennant (1975) obtained the best results when counts of one were given to a root crossing a line, a root ending touching a line and a curved portion touching a line. Counts of two were allocated to curved portions which lay on or along a line. However in these systems roots were floating on top of the counting lines. In our system, the counting lines are on top of the roots and adjacent grid elements are sharing counting lines. To make counting of intersections faster and more unambiguous, e.g. avoiding counting a single root crossing twice, we counted an intersection if a root touched or crossed the grid-line from above or from the left side. Roots intersecting, but not crossing, lines from below or from the right side were not recorded.

### Photography of tube rhizotrons

After shoot harvest the lower 0.5 m subsoil tube rhizotrons were removed and photographed. To avoid reflections from the PVC tubes they were removed (made possible by the lengthwise split) allowing the bare soil column to be photographed. The front and back side of tube rhizotron were photographed with a professional diffuse light source using a Canon EOS 60D camera with focal length of 83 mm and a fixed distance of 1 meter from the tube rhizotron.

### Root washing and root density measurements

After photography, the central 100 mm section of the lower 0.5 m subsoil soil columns, equivalent to the 1.70 to 1.80 m depth section, were separated for root washing. The roots were washed carefully from the soil using 1 mm sieves and stored in 50 % ethanol prior to cleaning for further measurement. The roots were placed on a tray (200 × 350 mm) and scanned using Epson Perfection V700 Photo. Root length measurements were obtained using WinRhizo® software (Regent Instruments Canada Inc., 2009).

In the software ImageJ (Rasband, 2014) the pictures were cut into sections showing the central 100 mm section of the 0.5 m core in the vertical direction, equivalent to the 1.70 to 1.80 m depth section. The pictures were then unwrapped using the software RoboRealm (Gentner, 2014) with the function Transform (Bottle_unwrap, radius set at 557). The pictures were cut to show the middle 80 mm of the tube rhizotron in the horizontal direction to avoid the most stretched vertical edges of the pictures. In ImageJ the height of the pictures were adjusted back to 100 mm since the RoboRealm Bottle_unwrap function had reduced the height. The resolution of the pictures was 1276 × 1598 pixels,and the picture size 80 mm × 100 mm corresponding to a pixel size of 63 µm.

The pictures of the tube rhizotrons were shown on a monitor in 2 × magnification. A grid printed on overhead plastic sheet was superimposed on the monitor (Fig. 1a). Horizontal and vertical root intersections were recorded for each individual grid element, using the procedure described for root intensity measurements on the rhizotron tubes. The superimposed grid on the monitor had a size of 160 × 200 mm with individual grid elements of 10 × 10 mm. This corresponded to an actual grid of 80 × 100 mm with individual grid elements of 5 × 5 mm. Thus each tube rhizotron was analyzed with a grid area of 160 cm2 and a total counting line length of 6.4 meter.

### Approaches to reduce the length of counting line

The root recordings were used for analyzing different approaches to reduce the length of counting line. This was done by systematically removing grid elements and repeating the analysis with the new grid pattern. The length of counting line was reduced from 6.4 m for each tube rhizotron to 3.2, 1.6, 0.8, 0.4 and 0.2 m in all three grid patterns (Fig 1). In the first pattern (design A), the sub-grid size was maintained and the grid area reduced from 160 cm2 (6.4 m of counting line) to 5 cm2 (0.2 m of counting line). In this design only the front side of the tube rhizotrons was analyzed when counting line was less than 6.4 m. In the second pattern, design B, the sub-grid size was increased while the grid area was maintained. In the third pattern, design C, the number of small grid line pieces (10 mm each) was decreased from 640 (6.4 m of counting line) to 20 (0.2 m of counting line) and the grid area maintained. In design B and C observations from both front and back sides of the tube rhizotrons were included for all lengths of counting line.

### Rooting depth measurements

On 2 July, rooting depth was measured on individual tube rhizotrons by recording the depth of the deepest root seen through the transparent interface on the front side of the tube rhizotron. Likewise, the depths below which 5, 10 and 25 roots were observed were also recorded on individual tube rhizotrons.

### Statistical Analysis

To quantify the variance of data obtained when applying different designs, coefficient of variation (CV) was calculated for each combination of grid design and length of counting line. This was done by fitting a model with cultivar and N treatments as factors in the GLM (General Linear Model) procedure of the SAS statistical package (SAS Institute Inc.). Cultivar and N treatment effects on rooting depth and on root intensity were calculated by analysis of variance using the GLM procedure of the SAS statistical package (SAS Institute Inc.).

### Results

Root intensity on the surface of the tube rhizotrons was positively and linearly correlated (p < 0.001) with RLD inside the tube rhizotrons (Fig. 2). We observed that root densities above ≈ 3 cm root cm-3 soil corresponded to high variation in root intensities (from 20 to 100 root intersections per m counting line). Therefore the correlation at these higher root densities appears to be weaker. At root densities below 1 cm root cm-3 soil few rhizotrons had roots recorded on the rhizotron surface. In several tube rhizotrons that contained less than 1 cm root cm-3 soil, no roots were recorded on the rhizotron surface.

**Figure 2:**
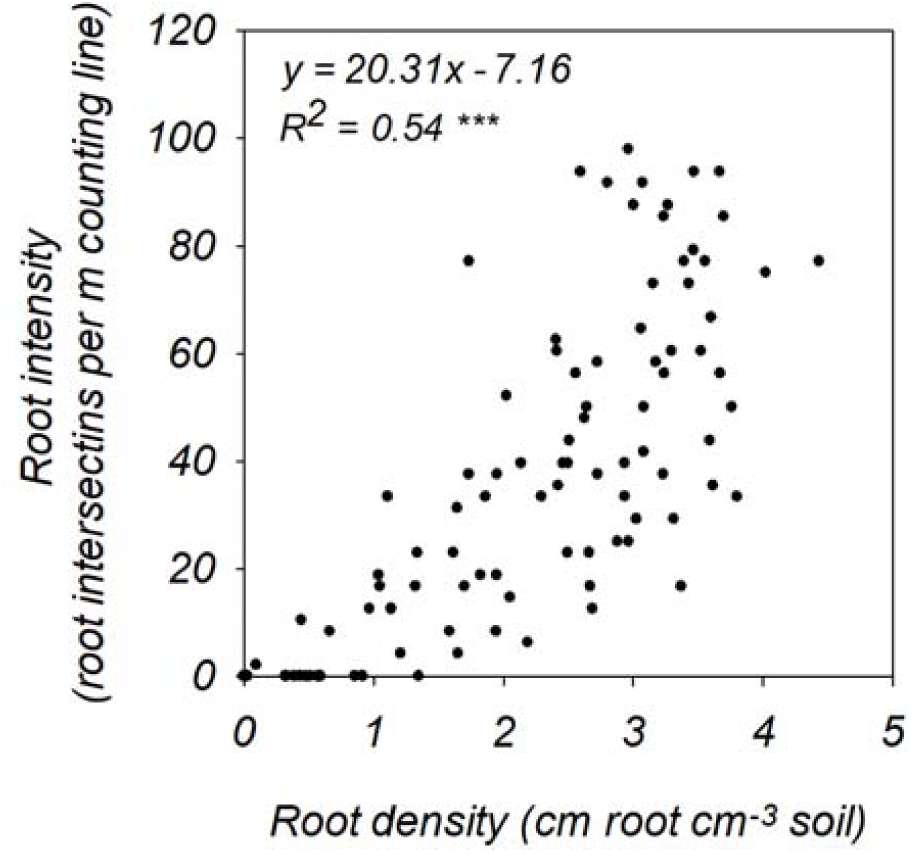
Root density in relation to root intensity of wheat roots growing in tube rhizotrons at rooting depths of 1.6 – 1.8 m. A linear regressing has been fitted to the plot. Root density measured by washing and analyzing roots in WinRhizo. Root intensity measured by superimposing a grid on the transparent tube rhizotrons and counting root and line intersections. The dimension of the grids was 60 × 80 mm with grid elements of 20 × 20 mm. Thus the length of counting line was 0.48 m for each grid.

### Effects of grid design on the variance of root intensity data

Fewer root-line intersections per grid were observed when counting lines were reduced (Fig.3a). Reduced length of counting line led to an increased variance of sampled data for all grid designs (Fig. 3b). This tendency was especially pronounced for the grid design A (Fig. 1a). Using design A the CV increased by approximately 10% every time the length of counting line was halved. By distributing the grid elements equally across the measuring area this effect could be minimized, as the variance remained low when this approach was used. In grid designs B and C (Fig. 1b and 1c) the counting line length could be reduced from 6.4 to 0.8 m without notable increases in the CV (Fig 3b). At counting line lengths above 0.8 m per grid, the designs B and C provided similar CV values of around 40 %. When length of counting line per grid was 0.4 m or less, design C was slightly better than design B as it gave lower CV values (Fig.3b).

**Figure 3:**
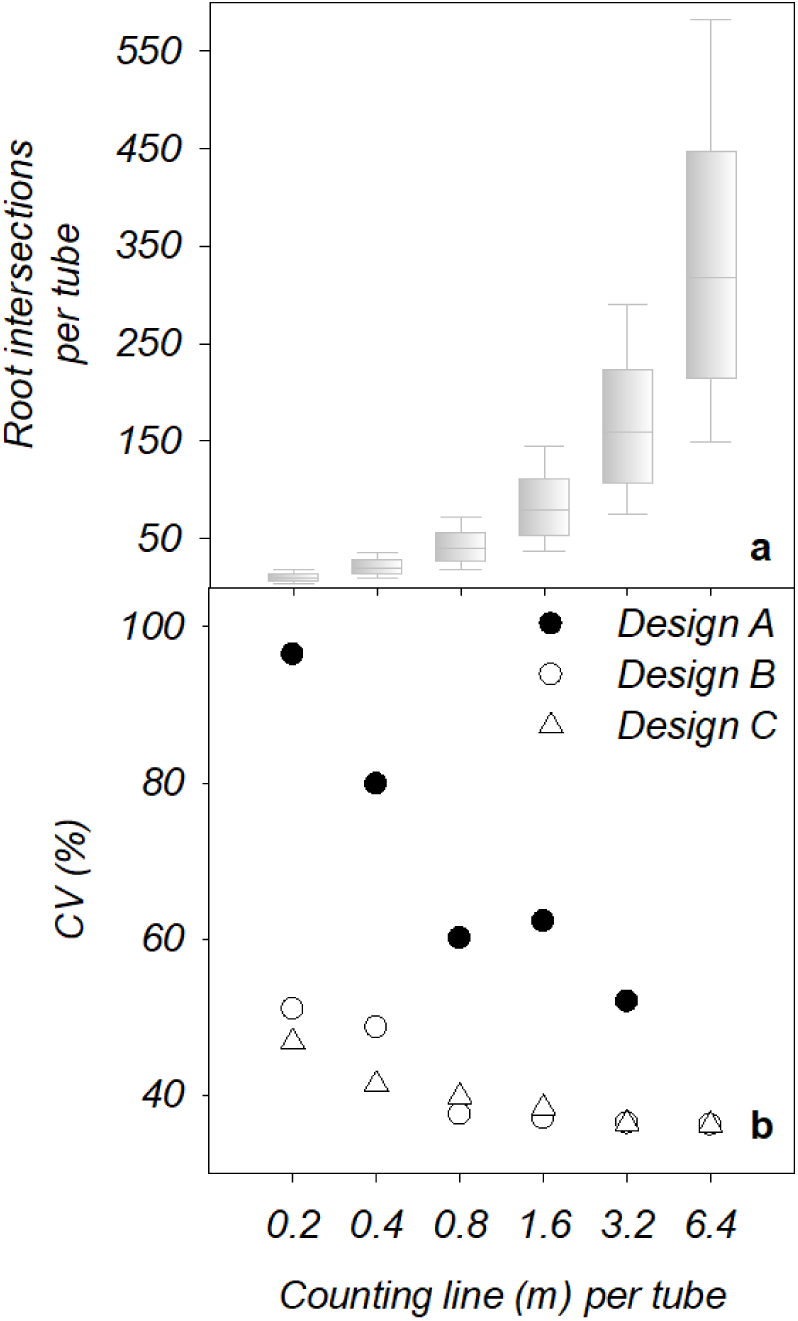
Root intensity measured by superimposing a grid on tube rhizotrons followed by counting the intersections with roots and lines. (a) Box plot of root intersections per grid when counting lines per grid increase. (b) Effects of different approaches to increase counting line length on the coefficient of variation (see Fig. 1). Coefficient of variation from ANOVA model using cultivars and nitrogen treatments as factors. Coefficient of variation when (•) maintaining size of grid elements and decreasing grid area, (◯) increasing the size of grid elements and maintaining grid area, (Δ) decreasing number of small line pieces and maintaining grid area.

### Effects of root intersections per grid on variance of root intensity data

The variance of sampled data increased when root-line intersections per grid decreased (Fig. 3a). For design B and C, CV increased when less than 50 root-line intersections per grid were recorded. This happened when the counting line was less than 0.8 m per grid. For design A, the number of root-line intersections per grid was dramatically higher – between 150 and 550 per grid at the same CV as seen in design B and C with 50 root line intersections per grid.

### Effect of cultivar on root intensity

The analysis of cultivar differences revealed that the orientation of observation lines seemed to influence whether cultivar effects could be detected. When observations of root intersections were limited to only include horizontal lines, the cultivars had significantly different root intensities (p = 0.03). No cultivar effect was seen when root intersections were counted only on vertical lines (p = 0.23) (grid size 2 × 80 cm2 and 3.2 m counting line) and when root intersections were counted on both horizontal and vertical lines (p = 0.10) (grid size 2 × 80 cm2 and 6.4 m counting line).

### Effects of different approaches to measure rooting depth

The estimated rooting depth decreased when the pre-determined number of deep roots counted to establish the depth was increased (Fig. 4). The different approaches to estimate rooting depth also affected the variance of the data as well as the differences between means. Using the depth of the deepest root only gave the highest variance of data (CV = 5.3 %). Increasing the number of roots associated with the estimated rooting depth reduced the variance of data and increased the difference between the sample means of the different cultivars. Consequently, the effect of cultivar on rooting depth was only significant when recording the soil depth below which 25 roots were observed (p-value 0.01). In contrast, recording the depth of the deepest root only resulted in a p-value of 0.65, while depths below which 5 and 10 roots were observed resulted in p-value of 0.23 and 0.14 respectively. Importantly, the ranking of the cultivars was also affected by the method of recording root depth. The cultivar that tended to have the deepest roots, when the estimate was based only on the deepest single root, was categorized among the more shallow rooted cultivars when applying recordings of the depth below which 5 or more roots were observed.

**Figure 4:**
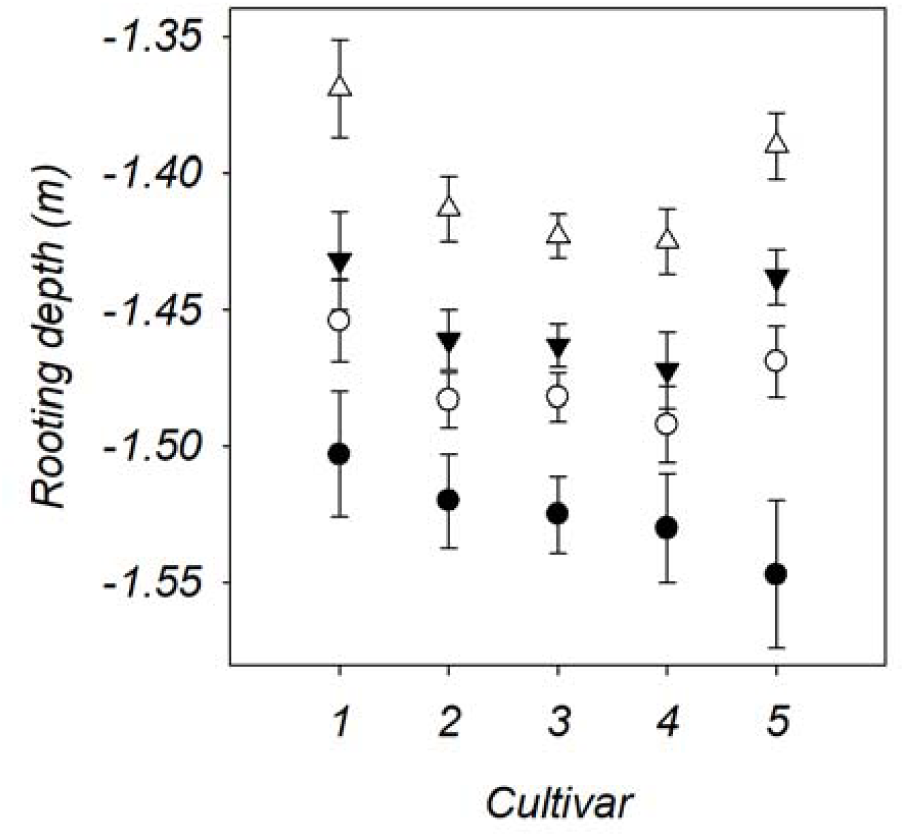
Mean rooting depths and S.E. measured with different approaches on 5 different wheat cultivars grown in tube rhizotrons. •The depth of the deepest root (CV = 5.3 %), ◯ the depth below which 5 roots (CV = 3.4 %), the depth below which ▾ 10 roots (CV = 3.5 %) and △ the depth below which 25 roots (CV = 3.7 %) are observed though the transparent tube rhizotrons.

## Discussion

The line intersect method is a simple and low-cost method allowing for detection of treatment and cultivar differences in rooting densities. In this study we investigated the importance of grid size and design, and of the relationship between root observations on the grids and root density determined by washing roots from the soil. The results showed that the relationship between root observations on the tube and RLD in the soil was best when root densities were less than ≈ 3 cm cm-3. In this study, root densities of less than 1 cm root per cm3 soil seemed to be the critical density below which roots on the tube-soil interface were often not observed despite their presence in washed samples. This threshold would be expected to decrease by the use of more appropriate grids. The grid used for correlation between root intensity and density only covered 60 out of 252 mm of the tube circumference. As the soil column was removed from the tube, roots were in some cases observed on other parts of the circumference, and frequently roots were observed to have grown in the lengthwise junction of the two halved tube pieces. The arrangement of the rhizotron tubes in this study made it difficult to view the “back” of the tubes, as they had to be lifted and rotated to do so. To facilitate easy access to the whole circumference of the rhizotron tubes the experimental design can be improved e.g. by place them on rotatable bases or suspend them from a frame.

The problem with detecting roots at low root intensities in rhizotrons and minirhizotrons can also be addressed by increasing the observation area per volume of soil. In minirhizotron studies this can be done by installing a higher number of minirhizotrons. In rhizotron studies the use of flat rhizotrons (Dresboll et al., 2013) increases the observation area per soil volume. However these solutions also increase the restriction of root growth and may potentially bias the results compared to a more natural three dimensional growth possible in circular tubes.

Another important observation from this study was that root intersections with vertical lines showed cultivar differences whereas intersections with horizontal lines did not. As lateral roots mainly have a horizontal growth direction they will dominate the intersection counted on vertical lines whereas main roots presumably dominate root intersections on horizontal lines. That differences between the cultivars were observed using vertical counting lines suggest that the cultivars may differ in their capacity for lateral root growth.

### Effects of grid design on the variance of root intensity data

The grid designs chosen in this study all used a systematic sampling strategy which has been shown to be more precise than random sampling strategies for these types of studies (Wolter, 1984; Wang and Qi, 1998). The grid design A (Fig. 1a) used a strategy of decreased measuring area for each soil section. This has the practical advantage of reducing the time needed both for image acquisition and time spent on counting roots. However, the results from the present study demonstrate that a grid design with reduced measuring area come at the cost of a high data variance. A strong reduction in variance of data down to CV values of around 40 % were obtained from grid designs with the counting line distributed on the entire soil section area. Reduction in measurement effort was made by reducing the line length in a way where the entire area was still included in the measurement. A CV around 40% is comparable to, or lower than what is normally obtained by other root measurement methods. With four replicates using the core break method or root washing of cores from field soil, CV values in the range from 37 to 71 % are normal (Kücke et al., 1995).

Increasing the length of counting line in specific soil layers will increase the number of observed line intersections. In this study, a counting line of 6.4 m per soil section resulted in 150 to 570 line intersections per soil section per tube. When grid designs B and C, in which the counting line is equally distributed across the entire measuring area was applied, the counting line could be reduced to 0.8 m resulting in 20 to 70 line intersections per soil section per tube without increasing the variability of data.

Based on the results obtained, we propose that the optimal length of counting line, when distributed optimally across the entire soil section area, can be defined based on the number of root-line intersections observed. For the tested data, a grid design resulting in 50 line intersections per soil section per tube ensured lowest data variability.

To the authors’ knowledge, this sampling approach, in which the number of positive observations determines the length of counting line, has not been described before. Other research fields have the same challenges as root research in gaining as many observations as needed to achieve adequate precision but no more, to avoid using unnecessary time and resources. However we can find only one previous study using a similar approach by defining the minimum number of positive observations needed for minimum variance. In wildlife research, Seaman et al., (1999) reported that variance approached an asymptote at about 50 observations per animal per home range.

The minimum number of observations needed can be expected to increase with an increase in the standard deviation. For uniformly repacked soil with the same number of replicates, as used in this study, the standard deviation for root observations can be expected to be in the same range as observed in this study. However in minirhizotron field studies, the soil is structured and less uniform, larger standard deviation can therefore be expected for these root observations. This approach, where 50 root-line intersections per soil section per tube per is aimed for, will be difficult to obtain at low root intensities. Alternatively more replicate rhizotrons may be needed to handle the higher variability at low root densities. It should be kept in mind though, that root observations are dynamic. In a soil layer where crop roots have just entered, and the density is still very low, the root density will be rising, and a few days later, the root density may be high enough to allow observation of 50 root intersections and thereby to get an estimate with a lower CV from the soil layer.

### Different approaches to measure rooting depth

Rooting depth, and the development of increased rooting depth during crop growth, has also been shown to be an important measure. Measurements, using more than just the single deepest root in each rhizotron showed to be the best approach due to larger differences in means between cultivars as well as a lower variance (the estimates were based on 5, 10 or 25 roots, rather than one). Thorup-Kristensen (1998) used measurements of the depth below which 3 roots were observed with success as he found significant differences (p ≤ 0.05) between 12 pea genotypes in field soil with four plot replicates and two minirhizotrons per plot. Studies where the depth of the deepest root has been used as a measure of rooting depth have been shown less effective in detecting genotypic differences in wheat (Wasson et al., 2014; Ytting et al., 2014; Rasmussen, unpublished results 2014).

For root depth measurements in rhizotrons and minirhizotrons it is rarely the absolute deepest root that is observed, and the final value is an average across several observations. Therefore it is not the maximum rooting depth that is effectively measured but rather the root intensities at depth. As an increased numbers of roots counted, the measurement is increasingly reflecting the root intensity in depth rather than the maximum rooting depth.

One advantage when using a higher number of root counts for rooting depth measurements are that these results are more likely to reflect the effective “functional” rooting depth for processes such as water or nutrient uptake better, both of which require a minimum density for significant resource utilization by plants. The effective functional rooting depth can be defined as the depth where the locally available resource can be utilized within a limited time.

Detection of the effective functional rooting depth is inherently problematic as it depends upon which root function is of interest. In the deeper soil layers this will often be water or N uptake. The uptake of N and water in specific soil layers depend both on RLD in the soil layer and inflow rate per length of root, which again depend on N/water demand of the crop as well as availability in other soil layers (Asseng et al., 1998; Haberle et al., 2006). Also the temporal aspect is important because roots that have a long time to explore a soil layer can have an effective total uptake even at low RLD and inflow rates if given enough time. Moreover RLD in a given soil layer can increase rapidly, until the plant reaches the reproductive phase, whereas especially water uptake often takes more time to measure. Furthermore in field subsoils, many roots are found in pores and cracks, resulting in reduced root-soil contact. This has a negative effect on the inflow rate per length of root (White and Kirkegaard, 2010).

Due to these characteristics of root growth and function, the effective rooting depth and minimum RLD required for effective water and nutrient uptake is not easily determined. For N uptake, the average maximum rooting depth, measured as the average depth of the deepest roots observed in minirhizotrons, has been shown to correlate well with the depth of functional N uptake by catch crops (Thorup-Kristensen, 2001). For wheat, the proportion of utilized N from depth in the period from tillering to flowering has been shown to correlate with rooting depth measurements at flowering but utilization in depth was reduced at low RLD (≈ 0.1 cm cm^−3^) in the deepest soil layers and available N in top-soil layers (Kuhlmann et al., 1989). For effective water uptake in wheat the results are more variable. Here a minimum RLD rather than the root depth that have been reported to govern water utilization. The minimum RLD necessary have been reported to be as low as 0.1 and up to 4 cm cm^−3^ for effective water uptake (Asseng et al., 1998; Barraclough et al., 1989; Xue et al., 2003; Kirkegaard et al., 2007). This was measured from anthesis to maturity by Kirkegaard et al. (2007) but the exact time period and date for measurements were not systematically reported by Barraclough et al. (1989), Asseng et al., (1998) and Xue et al. (2003).

Compared to measurements of root densities down the soil profile, measurements of the rooting depth are fast and easy do and to interpret as only one value per plant or plot is obtained. This study was performed on tube rhizotrons, but recordings of the depth below which a pre-determined number of roots are observed can easily be used in minirhizotron studies and core break studies as well.

One should keep in mind that often in root depth measurements it is fast measurements with low variance that are important as both contribute to make detection of genotypic and treatments differences possible. Using a higher number of roots counted increases the time required to make the measurements which make this approach less favorable. Also in recordings of very young plants with only few roots, the average depth of the deepest roots is likely to give more meaningful results.

## Conclusion

Improved root observation strategies can be developed for root observations on rhizotron and minirhizotron surfaces, leading to reduced statistical variance, while reducing the amount of work hours needed for observation.

Root intensity should be considered when deciding on a grid design in rhizotron studies using the line intersect method. When the root intensity is high, the length of grid line analyzed can be reduced, saving time for root counting at high root intensities. We found that a minimum of app. 50 root intersections per grid resulted in the lowest obtainable variance for grid patterns.

It is also important to distribute the observation area across as much of the rhizotron surface as possible, even when the grid line length analyzed is reduced. Much improved results were obtained when the counting grid was distributed across the entire rhizotron circumference, rather than on a selected area. For this study, a low CV was obtained with a minimum 0.8 m per rhizotron tube section, as long as the grid line was distributed around the whole circumference of the tube and a minimum of 50 roots were observed in the tube section.

Restricting the measuring line to a small sub-area of the tube rhizotron section drastically increased variance of data which makes this approach the least useful.

For root depth measurements, estimating root depth based on the 5, 10 or 25 deepest root observations reduced the observational variance as compared to observing the deepest root as has often been done previously. Measurements of the depth, below which 5, 10 or 25 roots were observed, gave lower variance and larger differences between means. Only by using a threshold of 25 roots were significant cultivar differences in rooting depths (p = 0.01) detected, as this measurement had the largest difference between means.

## Acknowledgements

We thank Anders Kristian Nørgaard and the rest of the staff at University of Copenhagen the Experimental farm in Taastrup. Founds were provided to Dr K. Thorup-Kristensen by University of Copenhagen, Department of Plant and Environmental Sciences.

